## Supplementary Information for "CoPheScan: phenome-wide association studies accounting for linkage disequilibrium"

#### Contents

**Supplementary Methods** **2**

**Supplementary Figures** **7**

**Supplementary References** **23**

### Supplementary Methods

##### Background: the coloc approach to colocalisation analysis

Coloc [1] begins by enumerating all possible events and their relative likelihoods. At any individual SNP, four mutually exclusive events are possible, with prior probabilities given by,

$$
\begin{aligned}
p_0 &: \text{SNP is not causally associated with either trait} \\
p_1 &: \text{SNP is causally associated only with trait 1} \\
p_2 &: \text{SNP is causally associated only with trait 2} \\
p_{12} &: \text{SNP is causally associated with both traits}
\end{aligned}
$$

where $p_0 + p_1 + p_2 + p_{12} = 1$.
While the user may set these parameters to values that reflect their prior beliefs, default values [2] are supplied $p_{12} = 5 \times 10^{-6}, p_1 = p_2 = 10^{-4}$. We call a configuration a set of events across all SNPs in a region. Subject to the constraint that at most one SNP is causally associated with each trait, each configuration corresponds to exactly one hypothesis:

$$
\begin{aligned}
H_0 &: \text{no association in the region} \\
H_1 &: \text{association to trait 1} \\
H_2 &: \text{association to trait 2} \\
H_3 &: \text{association to both traits, with different causal variants} \\
H_4 &: \text{association to both traits, with a shared causal variant}
\end{aligned}
$$

All possible configurations of the causal variants for each of the two traits are enumerated and weighted by their prior probabilities, and the support for each hypothesis is thus evaluated as a posterior probability.

##### Adaptation of coloc for a known causal variant

We consider the case where a SNP of interest is known to be causal for a phenotype (primary trait) which is often the case in PheWAS, and we are interested in determining if it is also causally associated with another phenotype (Fig.1a). We will refer to the variant of interest hereafter as the query variant and the phenotype tested as the query trait. At the query variant, exactly two events are possible with prior probabilities

$$
\begin{aligned}
p_c &: \text{query variant is also causally associated with the phenotype} \\
p_0^c &: \text{query variant is not causally associated with the phenotype}
\end{aligned}
$$

At the other variants, the events and their probabilities are

$$p_a : \text{non-query variant is causally associated with the phenotype}$$
$$p_0^a : \text{non-query variant is not causally associated with the phenotype}$$

Again, we can form configurations subject to the constraint of at most one causal variant. In a region with Q SNPs, each configuration belongs to one of three hypotheses:

$H_n$ : No association with the query trait (one configuration)

$H_a$ : Association of a variant other than the query variant with the query trait (Q-1 configurations)

$H_c$ : Association of the query variant with the query trait (one configuration)

**Adaptation of coloc priors for CoPheScan**

Under the assumption of at most one causal variant in a region for any trait, all or nearly all variants in any individual configuration will non-causal. For example, the prior probability of the configuration corresponding to $H_c$ is ${p_0^a}^{Q-1} p_c$. As the priors need to sum to 1 over the configurations compatible with the single causal variant assumption, the three prior probabilities corresponding to configurations in $H_n, H_a, H_c$, respectively, are proportional to $p_n, p_a, p_c$, subject to the constraint that $p_n + (Q-1)p_a + p_c = 1$. We assume $p_n \sim p_0^a \sim p_0^n$.
These can be either sampled using a hierarchical model or fixed values may be chosen *a priori* to reflect researcher belief. Given default values in coloc, these may be derived as

$$p_c = \frac{p_{12}}{p_{12} + p_1} \qquad\qquad p_a = \frac{p_2}{p_2 + p_0}$$

**Computation of posterior probabilities**

The posterior odds of each hypothesis for each configuration ($S$), given the data ($D$) for the query trait w.r.t the null hypothesis ($H_n$) is given by,

$$\frac{P(H|D)}{P(H_n|D)} = \frac{P(S)}{P(S_n)} \times \sum_{S \in H} \frac{P(D|S)}{P(D|S_n)} \tag{1}$$

where $D$ represents the observed data, $S$ represents any configuration compatible with the hypothesis of interest, $H$, and $S_n, H$ respectively represent the null configuration and hypothesis of no association at any SNP. In (1), the first ratio in the RHS is the prior odds and the second ratio is the Bayes Factor (BF). The posterior odds of each hypothesis is given by,

$$\begin{aligned}
\frac{P(H_n|D)}{P(H_n|D)} &= 1 \\
\frac{P(H_a|D)}{P(H_n|D)} &\simeq \frac{p_a}{p_n}\sum_{j\neq c} BF_j = \frac{p_a}{p_n}\tilde{BF}_a \\
\frac{P(H_c|D)}{P(H_n|D)} &\simeq \frac{p_c}{p_n} \times BF_c = \frac{p_c}{p_n}\tilde{BF}_c
\end{aligned} \tag{2}$$

where BF is the Bayes factor summarising evidence for association at SNP $j$. $\tilde{BF}$ and $BF$ refer to overall and per-SNP hypothesis Bayes factors respectively.

###### Bayes factors

CoPheScan can use one of two forms of Bayes factor:
**Wakefield's Approximate Bayes Factor** [3], is the relative support for a model where the SNP is associated with a trait compared to the null model of no association at that SNP. The above formulation takes no account of association at other SNPs, so its use holds under the single causal variant assumption, but not in cases where a given region has multiple causal variants.
Recently, the coloc approach was extended to deal with multiple causal variants by incorporating the **Sum of Single Effects (SuSiE)** Bayesian fine mapping regression framework. [4, 5]. We adapt the same approach to account for the multiple variants in the CoPheScan context. SuSIE takes in GWAS summary statistics and the linkage disequilibrium (LD) matrix as input and returns credible sets, each containing a set of strongly correlated SNPs that are associated with the given trait. One of the variants in each credible set has a high probability of having a non-zero effect. In the known causal variant case, we apply SuSiE on the queried trait and extract the vectors of Bayes factors (BF) returned for each of the regressions, and evaluate the posterior support for each hypothesis for each inferred SuSiE regression.

###### A hierarchical model to estimate priors with multiple phenotypes

As seen previously from the analysis of a single trait $k$, we can summarise the data with the quantities

$$\tilde{BF}_{ak} = \frac{P(D_k|H_{ak})}{P(D_k|H_{nk})} = \sum_{j\neq c} BF_{jk}$$

$$\tilde{BF}_{ck} = \frac{P(D_k|H_{ck})}{P(D_k|H_{nk})} = BF_{ck}$$

where we introduce subscript $k$ to indicate quantities related to trait $k$.
The likelihood of the data can be expressed as a product over the likelihoods of the data for each trait ($L_k$), assuming these data are independent,

$$L = \prod_k L_k$$

where the $L_k$, the likelihood for the data for trait $k$ only, can be expressed as,

$$\begin{aligned} L_k &= P(D_k|H_{nk})P(H_{nk}) + \sum_{S \in H_a k} P(D_k|S)P(S) + P(D_k|H_{ck})P(H_{ck}) \\ &= P(D_k|H_{nk})p_n + P(D_k|H_{ak})p_a + P(D_k|H_{ck})p_c \\ L_k &\propto p_n + \tilde{BF}_{ak}p_a + \tilde{BF}_{ck}p_c \end{aligned} \tag{3}$$

##### MCMC sampling from the posterior

We use Metropolis-Hastings to sample from the posterior distribution of $p_a, p_c$. We set

$$\frac{p_a}{p_n} = e^{\alpha} \qquad \frac{p_c}{p_n} = e^{\alpha+\beta} \tag{4}$$

where

$$\alpha \sim N(-10, 0.5) \qquad \beta \sim \Gamma(2, 0.5)$$

Thus the target posterior is

$$P(\alpha, \beta|D) \propto P(\alpha)P(\beta) \prod_k L_k \tag{5}$$

We set up a Metropolis-Hastings algorithm to sample from this distribution, at each step proposing a new $(\alpha, \beta)$ as

$$\alpha' \sim N(\alpha, 0.5) \qquad \beta' \sim \Gamma(\beta, 0.5)$$

and accepting the proposal if $P(\alpha', \beta'|D) > P(\alpha, \beta|D)$ or with probability $\frac{P(\alpha',\beta'|D)}{P(\alpha,\beta|D)}$ otherwise.
When measurements of genetic correlation $r_g$ between traits are available, we can adapt 4 as

$$\frac{p_{ak}}{p_{nk}} = e^{\alpha} \qquad \frac{p_{ck}}{p_{nk}} = e^{\alpha+\beta+\gamma r_g} \tag{6}$$

where

$$\begin{aligned} \alpha &\sim N(-10, 0.5) \\ \beta &\sim \Gamma(2, 0.5) \\ \gamma &\sim \Gamma(2, 0.5) \end{aligned}$$

proposing

$$\gamma' \sim \Gamma(\gamma, 0.5)$$

and accepting the proposal if $P(\alpha', \beta', \gamma'|D) > P(\alpha, \beta, \gamma|D)$ or with probability $\frac{P(\alpha',\beta',\gamma'|D)}{P(\alpha,\beta\gamma|D)}$ otherwise.

##### Computational time

CoPheScan is implemented in R and C++ and is available on CRAN from `https://cran.r-project.org/package=cophescan`.

The computational time of the MCMC model is dependent on the size of the variant and trait sets considered. For 100 variants and 1000 traits, 100,000 iterations takes 25 minutes on a single Intel Ice Lake CPU of a high-performance computing cluster. Timing scales linearly with the number of variant/trait pairs and the number of iterations as shown in Figure 1.

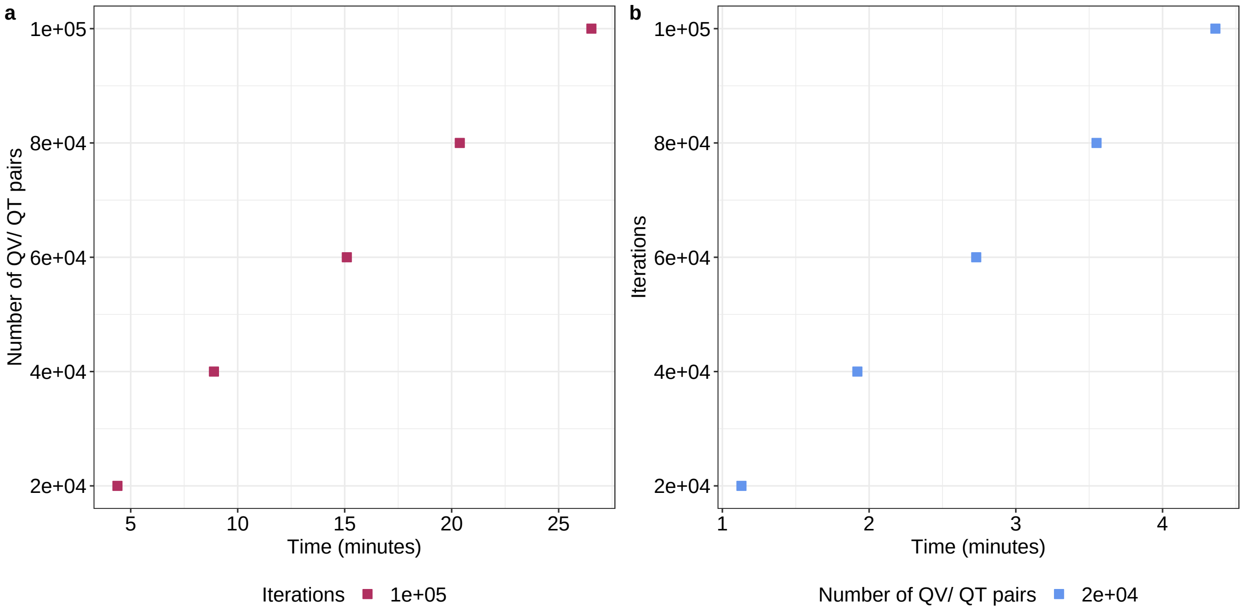


**Figure 1:** Computational time for the CoPheScan MCMC model in minutes (x axis) where the number of query variant - query trait (QV/QT) pairs (y axis) were varied and run for 100,000 iterations in (a) and the number of iterations (y axis) were varied and run with 100,000 QV/QT pairs in (b).

**Supplementary Figures**

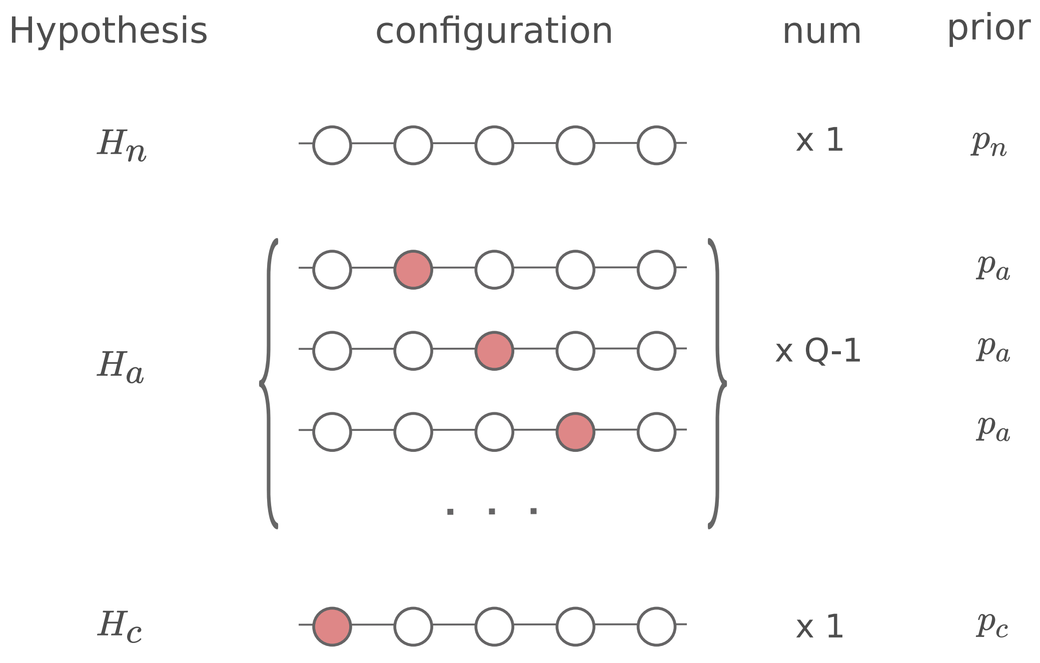


**Supplementary Figure 1:** The three CoPheScan hypotheses: Hn, Ha and Hc and their corresponding SNP configurations in a genomic region with Q SNPs (circles). The first SNP here is the query variant and the SNPs are shaded red when causal for the query trait.

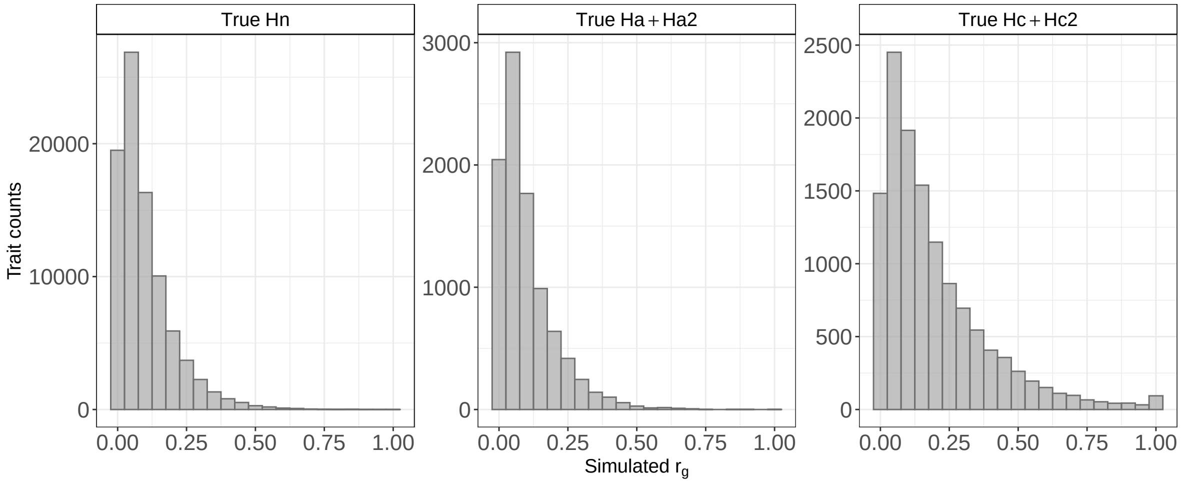


**Supplementary Figure 2:** Histograms of simulated genetic correlation ($r_g$) values simulated for the traits which are divided in panels based on the true hypothesis of the simulated traits.

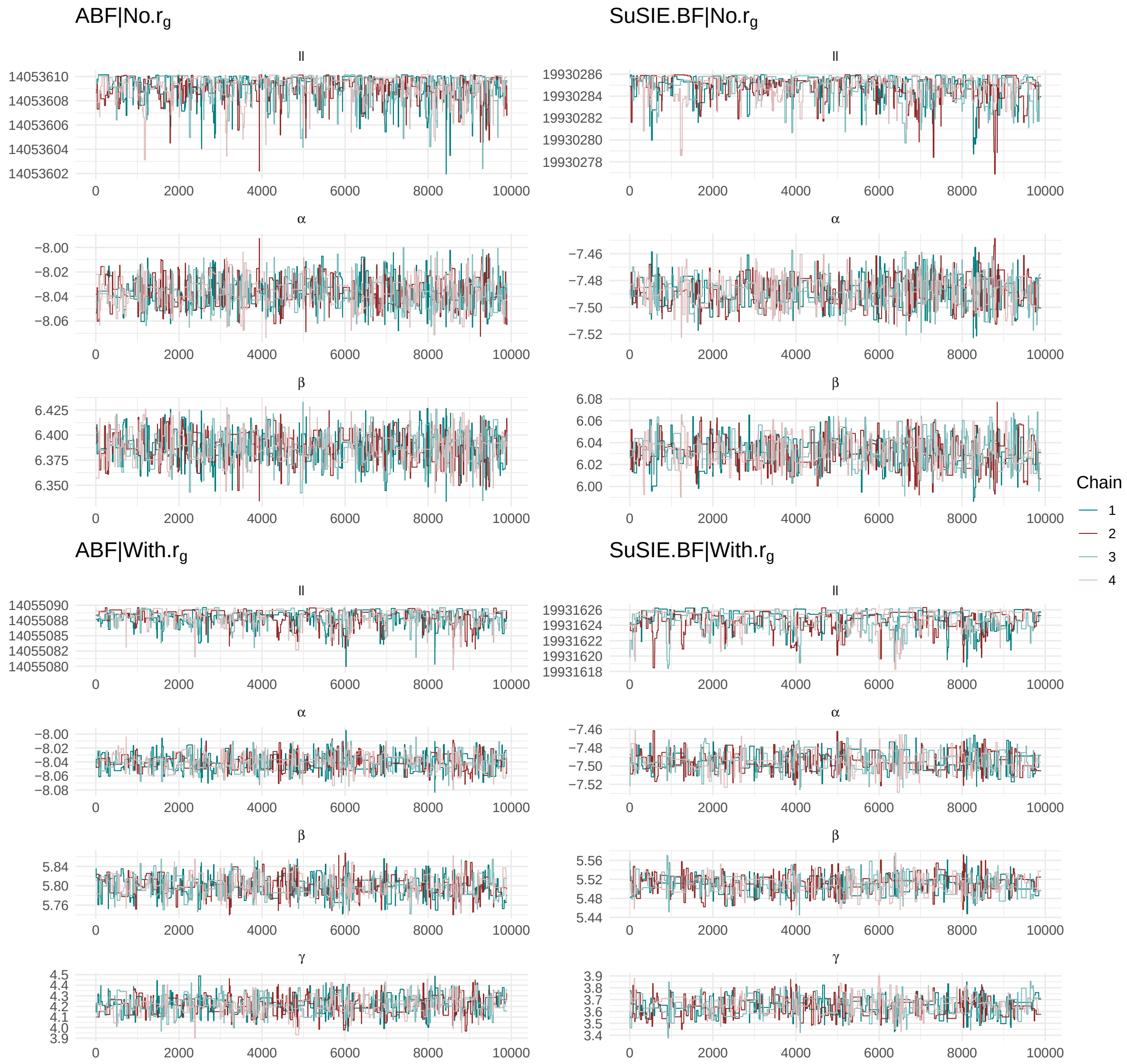


**Supplementary Figure 3:** Diagnostic trace plots of the hierarchical MCMC models on simulated data run with different approaches of CoPheScan. The models without $r_g$ were run for 3e5 iterations and those with $r_g$ for 1e6 iterations and the chains were thinned by retaining every 30th observation and 100th observation respectively. The trace plots are of the remaining 1e4 observations after trimming the first 100. (No $r_g$=3e3 burn-in, With $r_g$= 1e4 burn-in). ll=log likelihood.

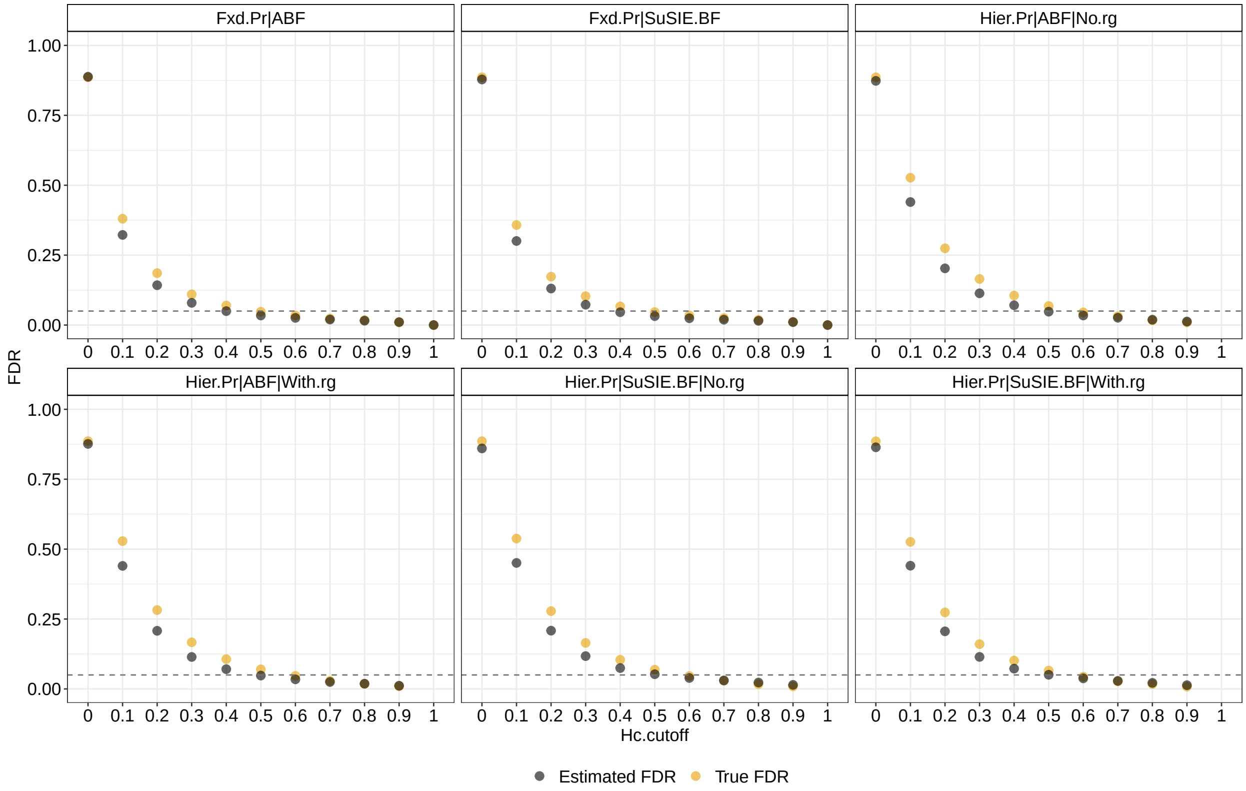


**Supplementary Figure 4:** False discovery rate (FDR) calculated for a range of [0-1] ppHc cutoff. True FDR is the FDR calculated using FP/(FP+TP) where FP and TP are the number of False and True positive cases identified using CoPheScan at each pp.Hc cutoff. Estimated FDR was obtained by calculating mean(1-ppHc) of the traits called as Hc by CoPheScan at the corresponding pp.Hc cutoff. The horizontal dashed line represents 0.05 FDR. .3

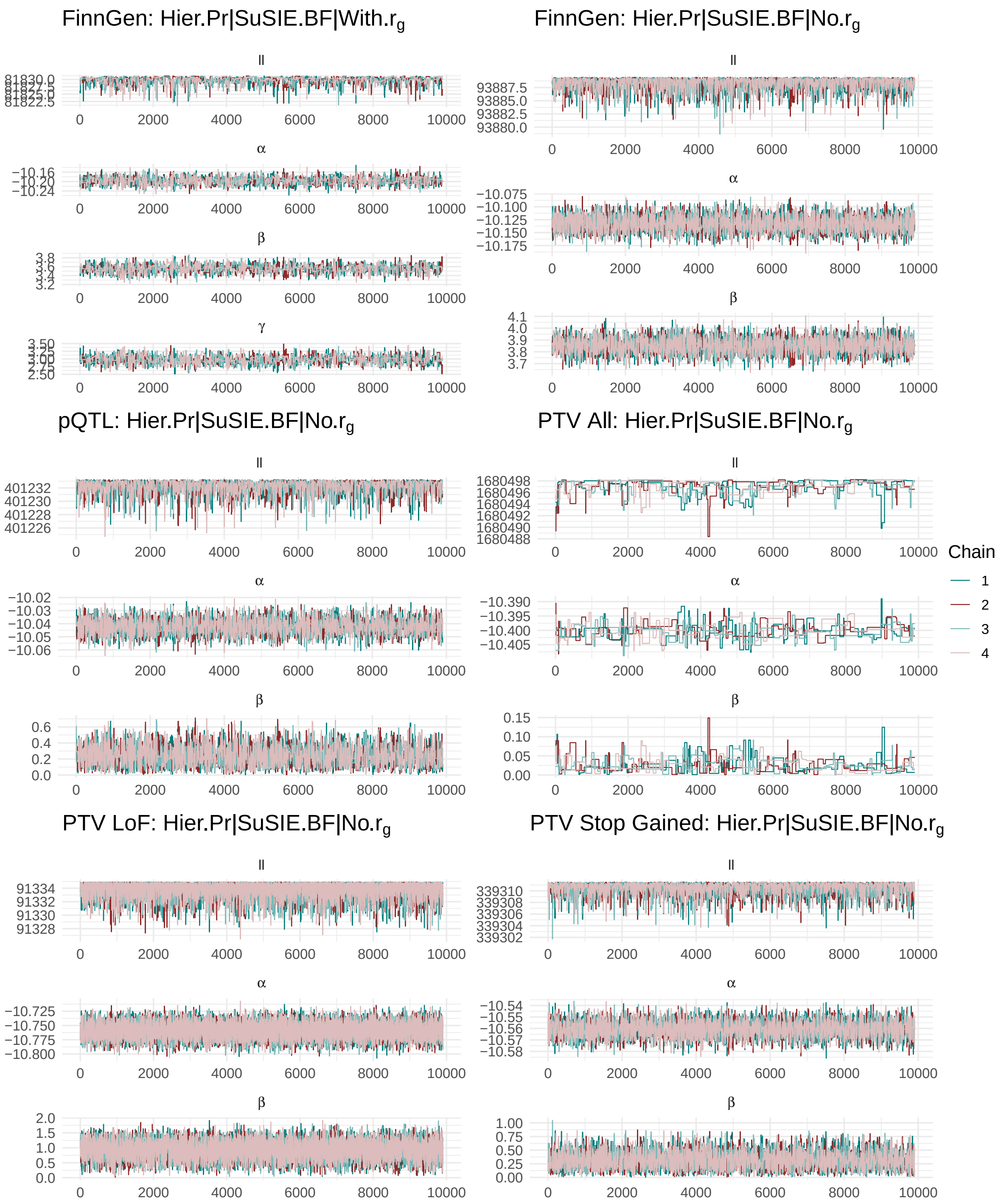


**Supplementary Figure 5:** Diagnostic trace plots of the hierarchical MCMC models on real data derived from the three different variant sources. The models were run for 3e5 iterations chains were thinned by retaining every 30th observation. The trace plots are of the remaining 1e4 observations after trimming the first 100. (3e3 burn-in). ll=log likelihood.

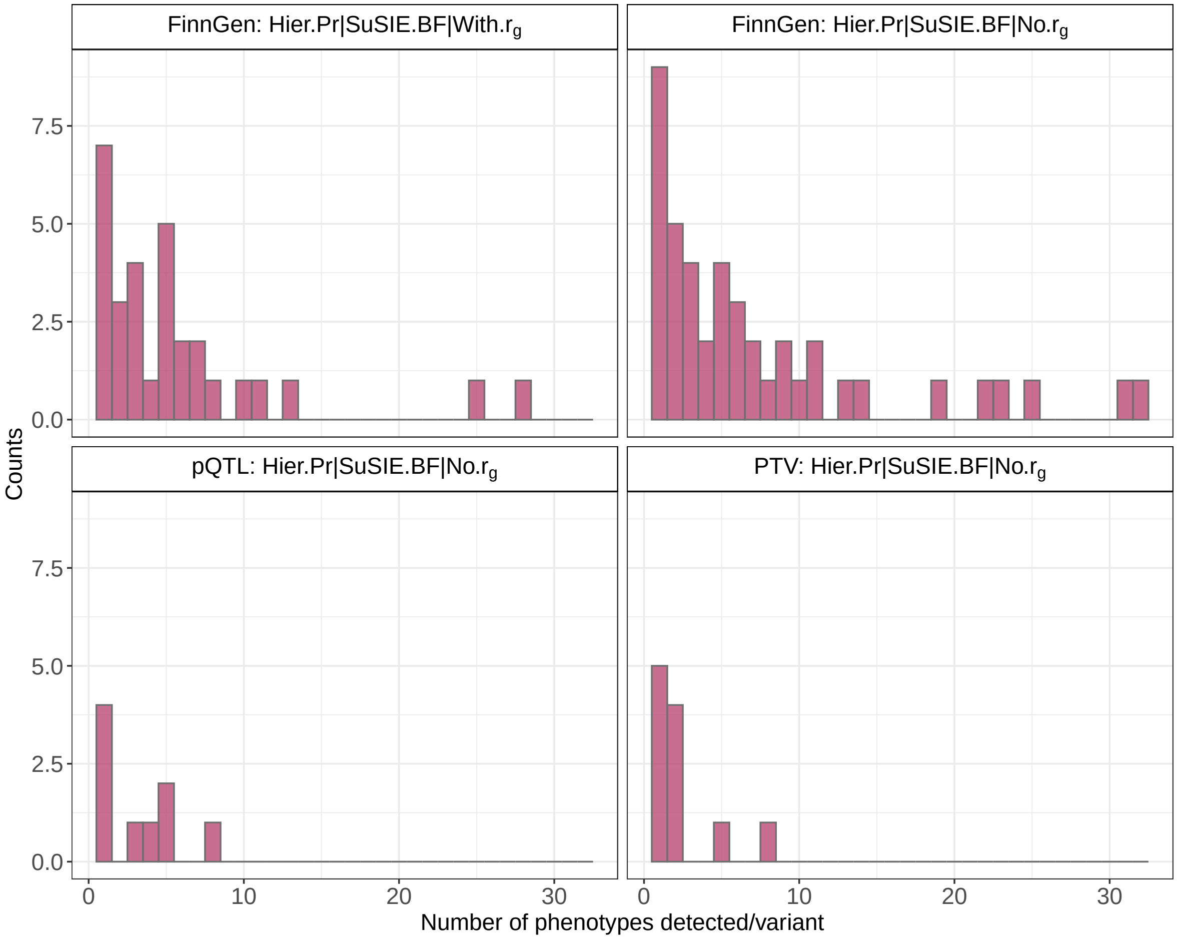


**Supplementary Figure 6:** Histograms showing the distribution of the number of phenotypes detected per variant with at least one variant detected.

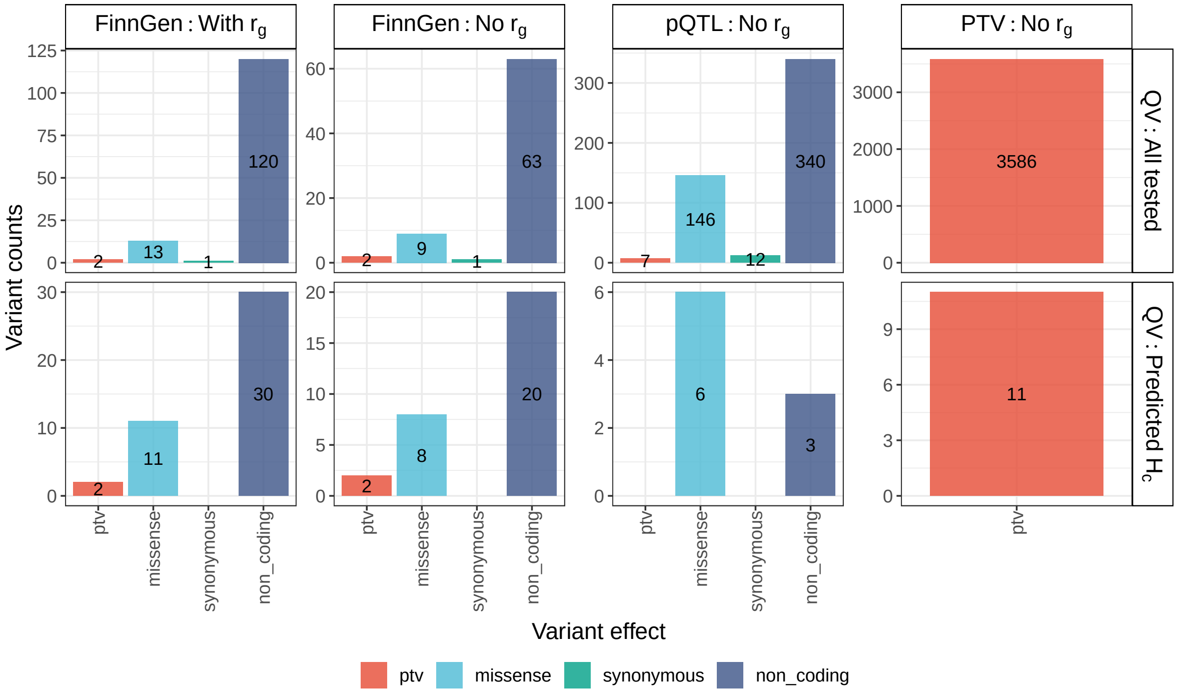


**Supplementary Figure 7:** Variant effects of all the query variants (QV) used in the study. The columns represent the sources of the QV. The first row shows the distribution of the variant effects of all QV used while the bottom row shows those of the ones detected as having a causal association with the UKBB traits (Predicted $H_c$).

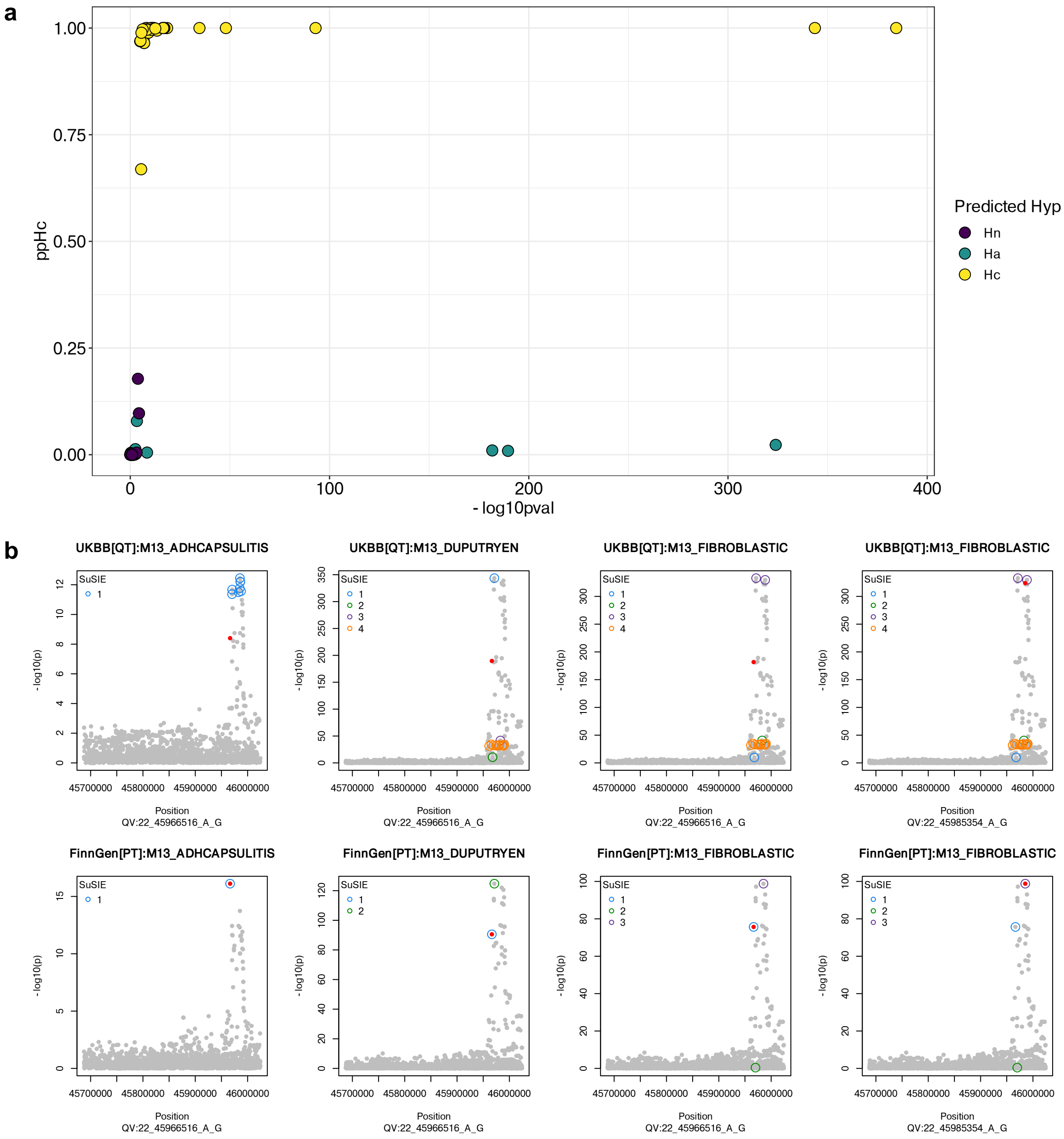


**Supplementary Figure 8:** (a) $ppH_c$ Vs. -log10(pval) of associations tested with CoPheScan for UKBB traits which were identical to those derived from FinnGen. The associations are coloured based on the predicted hypothesis. (b) Top row shows the Manhattan plots of four UKBB traits in the region around the query variants (red) whose -log10(pval) associations with traits were $> 5$ and were predicted as $H_a$. SuSIE credible sets identified are shown as colored circles around the SNPs. The bottom row shows the corresponding figures from the FinnGen dataset from which the variants identified by SuSIE were selected. Note: the region analysed with SuSIE for these traits in FinnGen (44.5MB - 47.5MB) was larger than the UKBB region (45.7MB - 46MB). QV=Query Variant, QT=Query Trait, PT=Primary Trait.

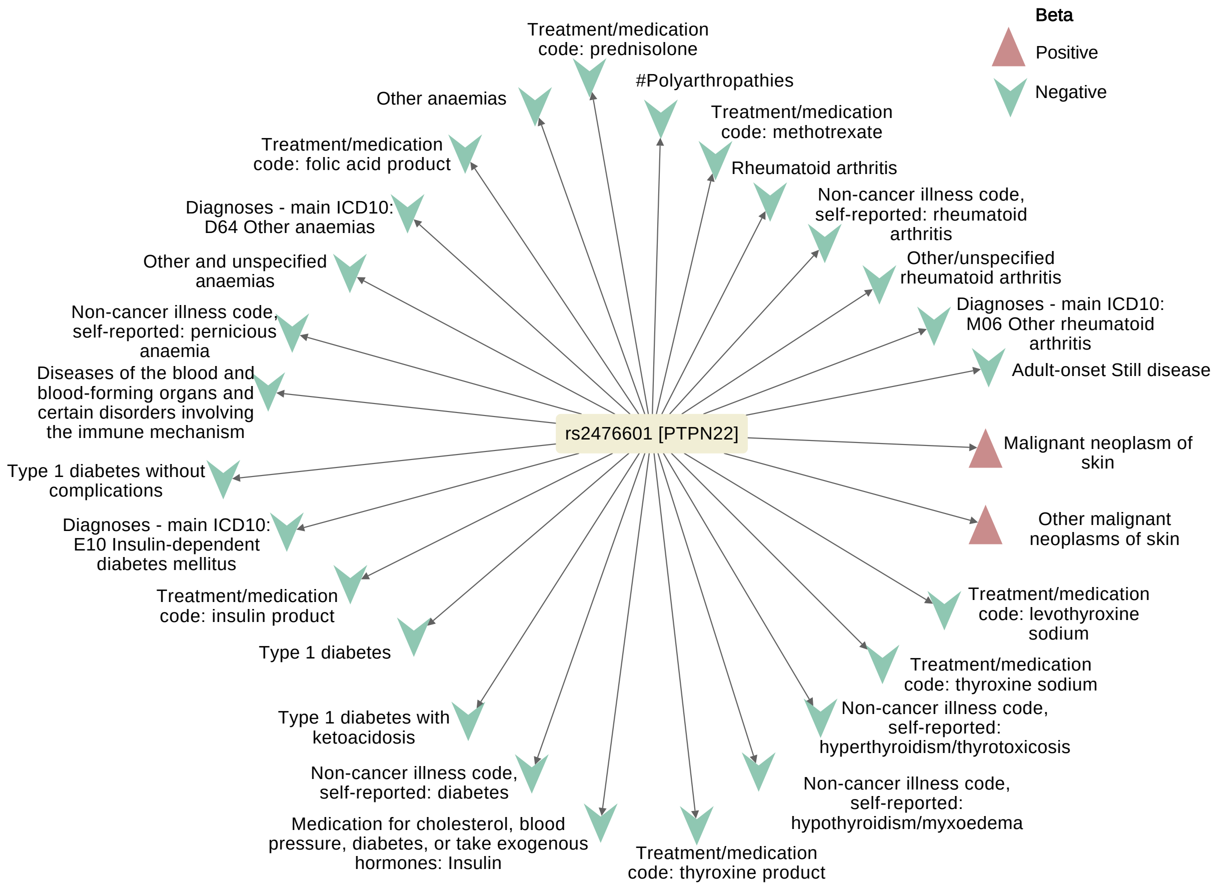


**Supplementary Figure 9:** Plot showing Hc associations (ppHc $>$ 0.56) of the *PTPN22* variant rs2476601_A>G. The direction of beta is shown with respect to the ALT allele (G).

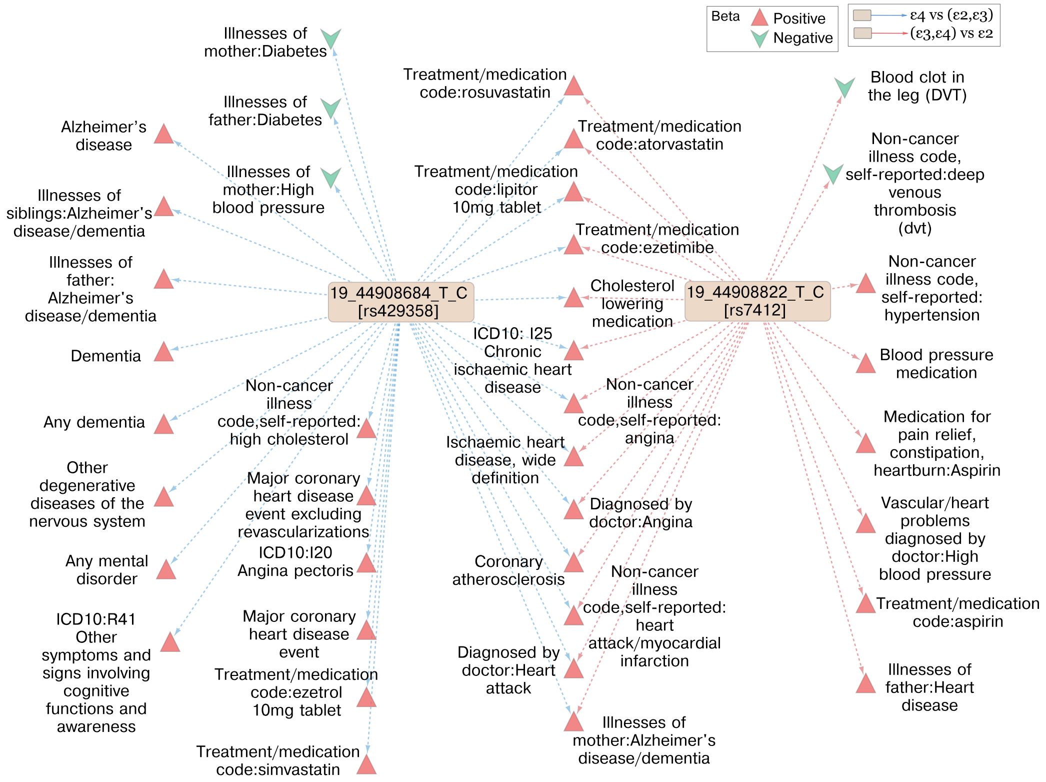


**Supplementary Figure 10:** Hc associations (ppHc $> 0.6$) of the APOE variants rs429358_T>C and rs7412_T>C. The direction of effect (beta) is shown with respect to the C allele at each SNP. The colour and shape of the trait nodes denote the direction of the estimated effect and the colour of the lines whether this relates to $\epsilon$4 compared to $\epsilon$2,$\epsilon$3 or $\epsilon$4,$\epsilon$3 compared to $\epsilon$2.

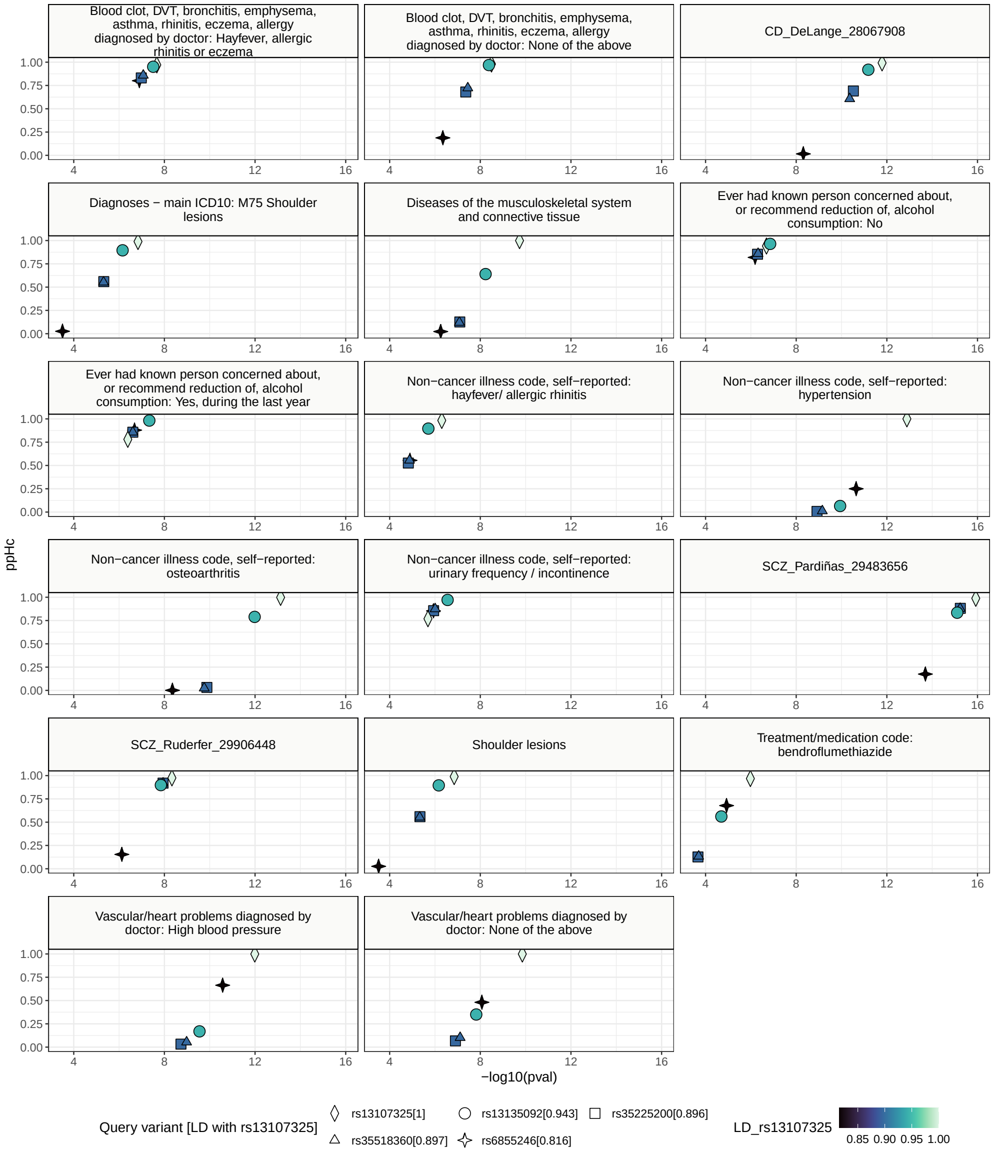


**Supplementary Figure 11:** $ppH_c$ Vs. -log10(pval) plots, from the sensitivity analysis of *SLC39A8* variants, of traits associated with rs13107325_C>T, a pleiotropic variant, and all variants in LD $>$ 0.8 with it.

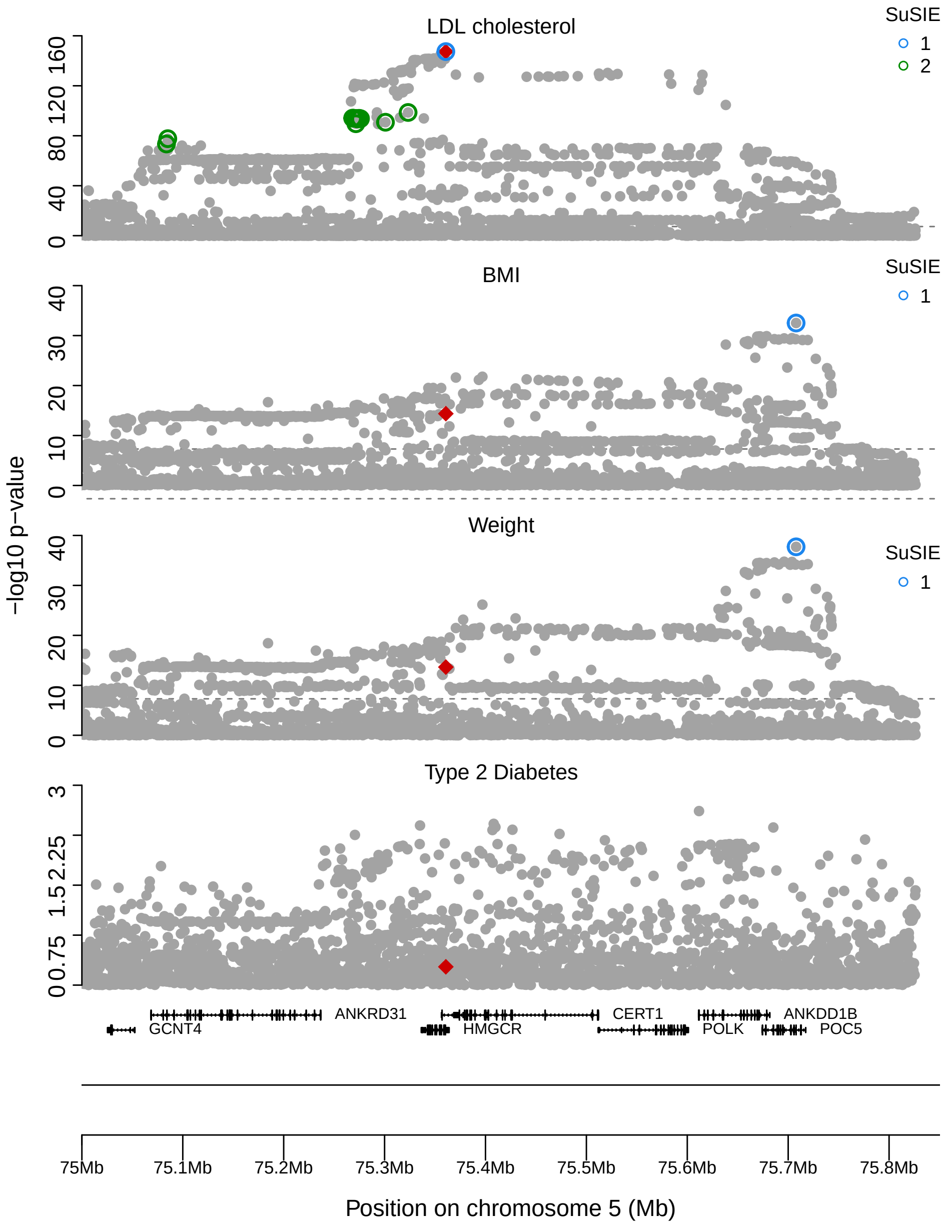


**Supplementary Figure 12:** Regional Manhattan plots of variants around the *HMGCR* variant, rs12196_T>C (red), in the UKBB traits: LDL cholesterol, BMI, weight and type 2 diabetes. SuSIE credible sets identified are shown as colored circles around the SNPs. rs12196_T>C is identified in a SuSIE credible set only in the LDL trait. Only one credible set was identified in the BMI and weight traits in the *POC5* gene region and none in type 2 diabetes.

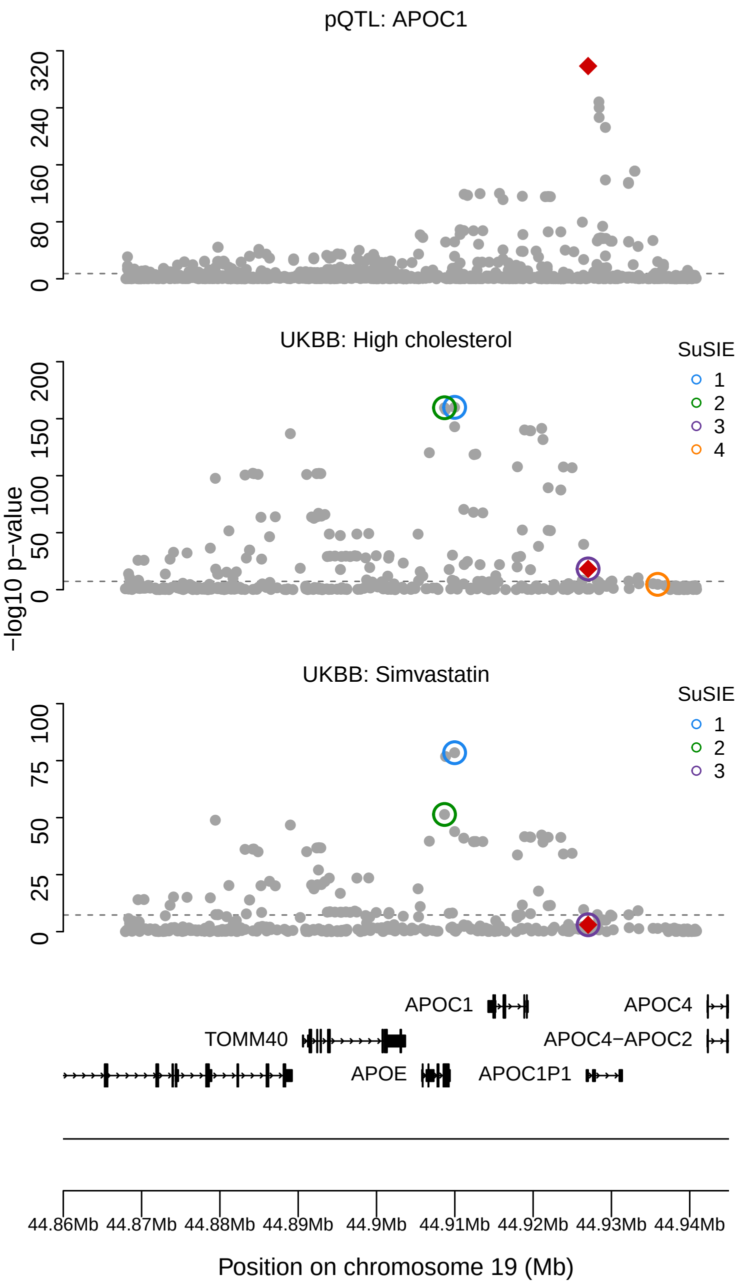


**Supplementary Figure 13:** Regional Manhattan plots of variants around the rs5112_C>G variant (red), a pQTL of *APOC1*, in the *APOC1*, High cholesterol (UKBB) and Simvastatin (UKBB) traits. SuSIE credible sets identified are shown as colored circles around the SNPs.rs5112_C>G is detected by SuSIE (purple circle) in both UKBB traits. The other two SNPs detected with SuSIE, rs429358_T>C and rs1065853_G>T (green and blue), are pQTLs of *APOE* [6] and might be independently affecting high cholesterol through a separate pathway.

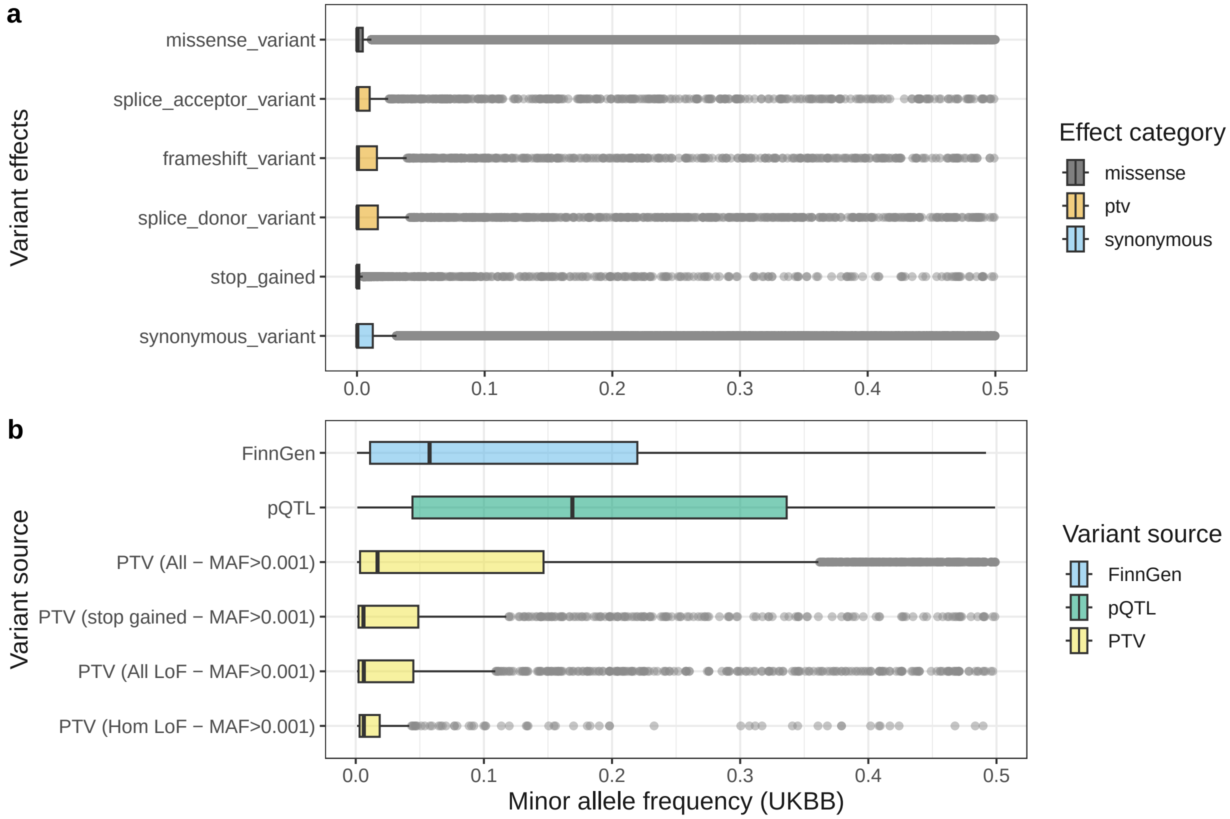


**Supplementary Figure 14:** Box plots showing the distribution of minor allele frequencies in a) UKBB variants grouped based on their effects b) All variants tested from different sources, which are filtered to exclude MAF <= 0.001.

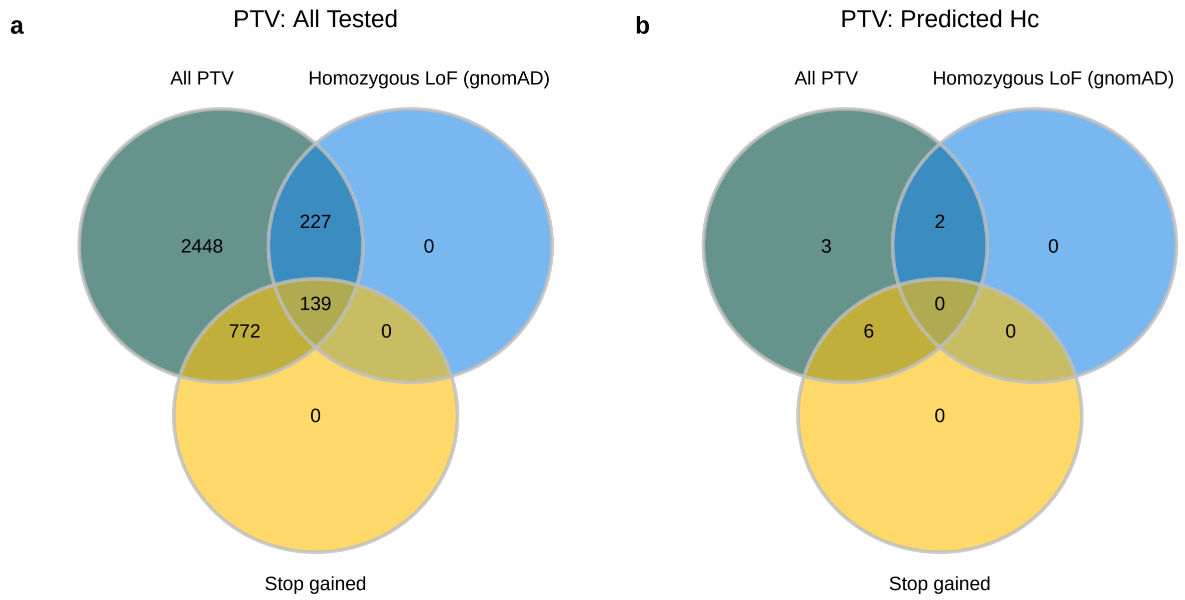


**Supplementary Figure 15:** Venn diagrams showing the protein-truncating variant (PTV) sets used in this study where a) shows all PTV used for testing and b) shows PTV predicted as Hc with UKBB traits. All PTV: UKBB PTV variants with MAF $>$ 0.001, Homozygous LoF (gnomAD): UKBB PTV (MAF $>$ 0.001) in common with the homozygous LoF from gnomAD v2.1.1 data set [7] and Stop gained: UKBB stop gained PTV (MAF $>$ 0.001)

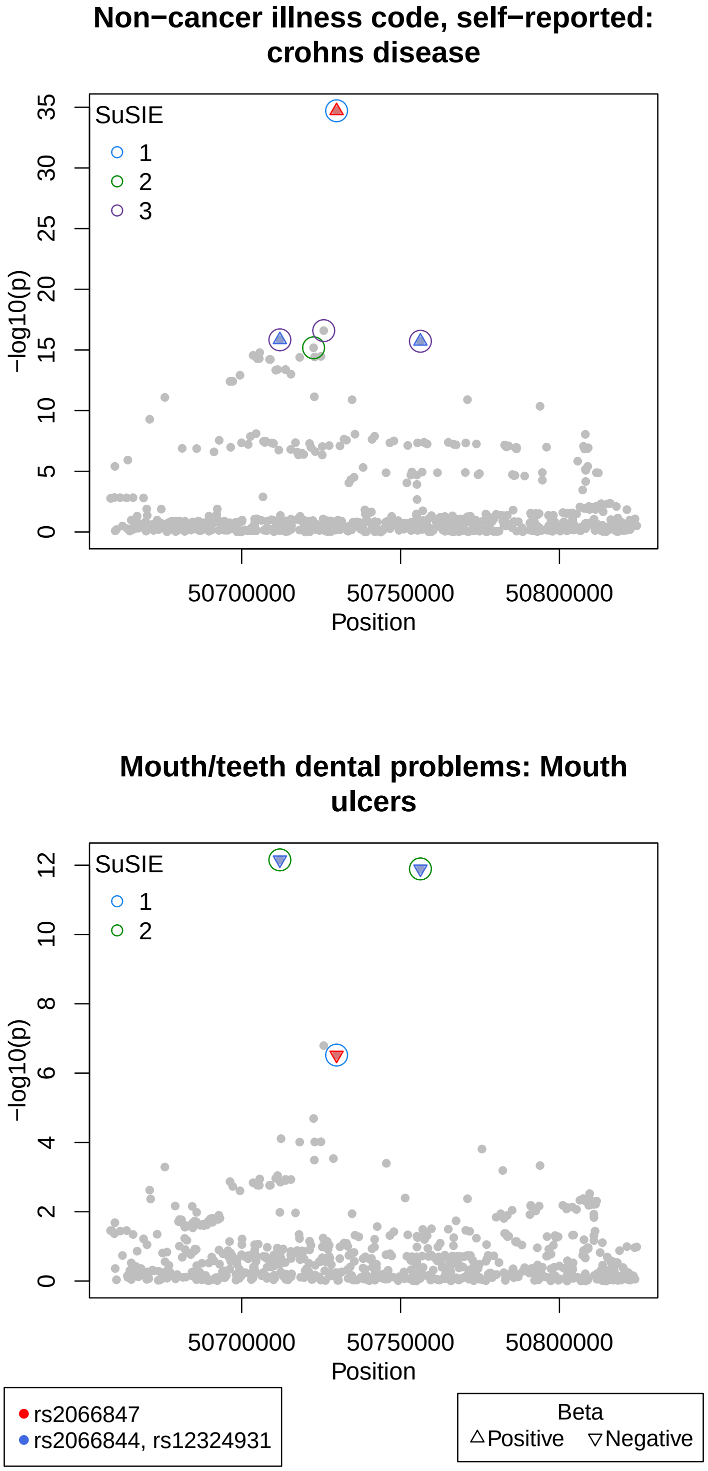


**Supplementary Figure 16:** Regional Manhattan plots of variants around the *NOD2* variant, rs2066847_G>GC (red), in the Crohn's disease (20002_1462) and mouth ulcer traits (6149_1). SuSIE credible sets identified are shown as colored circles around the SNPs. Two additional SNPs - rs2066844_C>T and rs12324931_A>C found by SuSIE in both traits are shown in blue.

### References


[1] Giambartolomei, C. *et al.* Bayesian test for colocalisation between pairs of genetic association studies using summary statistics. *PLoS Genet.* **10** (5), e1004383 (2014) .

[2] Wallace, C. Eliciting priors and relaxing the single causal variant assumption in colocalisation analyses. *PLoS Genet.* **16** (4), e1008720 (2020) .

[3] Wakefield, J. Bayes factors for genome-wide association studies: comparison with P-values. *Genet. Epidemiol.* **33** (1), 79–86 (2009) .

[4] Wallace, C. A more accurate method for colocalisation analysis allowing for multiple causal variants. *PLoS Genet.* **17** (9), e1009440 (2021) .

[5] Wang, G., Sarkar, A., Carbonetto, P. & Stephens, M. A simple new approach to variable selection in regression, with application to genetic fine mapping (2020). Issue: 5 Pages: 1273–1300 Publication Title: Journal of the Royal Statistical Society: Series B (Statistical Methodology) Volume: 82.

[6] Sun, B. B. *et al.* Genomic atlas of the human plasma proteome. *Nature* **558** (7708), 73–79 (2018). Number: 7708 Publisher: Nature Publishing Group .

[7] Karczewski, K. J. *et al.* The mutational constraint spectrum quantified from variation in 141,456 humans. *Nature* **581** (7809), 434–443 (2020). Number: 7809 Publisher: Nature Publishing Group .
